# Genetic control of transition from juvenile to mature wood with respect to microfibril angle (MFA) in Norway spruce (*Picea abies*) and lodgepole pine (*Pinus contorta*)

**DOI:** 10.1101/298117

**Authors:** Haleh Hayatgheibi, Nils Forsberg, Sven-Olof Lundqvist, Tommy Mörling, Ewa J. Mellerowicz, Bo Karlsson, Harry Wu, M Rosario García Gil

**Author notes:** Corresponding author: M Rosario García Gil.

## Abstract

Genetic control of microfibril angle (MFA) transition from juvenile to mature was evaluated in Norway spruce and lodgepole pine. Increment cores were collected at breast height from 5,618 trees in two 21-year-old Norway spruce progeny trials in southern Sweden, and from 823 trees in two 34-35 – year-old lodgepole pine progeny trials in northern Sweden. Radial variations in MFA from pith to bark were measured for each core using SilviScan. To estimate MFA transition from juvenile to mature, a threshold level of MFA 20° was considered and six different regression functions were fitted to the MFA profile of each tree after exclusion of outliers, following three steps. The narrow-sense heritability estimates (*h*^2^) obtained for MFA transition were highest based on the slope function, ranging from 0.21 to 0.23 for Norway spruce and from 0.34 to 0.53 for lodgepole pine, while *h*^2^ were mostly non-significant based on the logistic function, under all exclusion methods. Results of this study indicate that it is possible to select for an earlier MFA transition from juvenile to mature in Norway spruce and lodgepole pine selective breeding programs, as the genetic gains (*∆*_*G*_) obtained in direct selection of this trait were very high in both species.

## Introduction

Wood properties have become an important focus in advanced tree breeding programs, along with growth, vitality and form traits (Bouffier et al. 2009; Isik et al. 2008). The variability in wood properties within a tree is very large (Larson 1967), owing to the within-ring differences, the changes along a radius from pith to bark, and the differences associated with different heights in the tree (Zobel and Van Buijtenen 1989).

Differences between juvenile and mature wood are the major sources of variation in wood quality, both among and within trees (Zobel and Sprague, 1998). Such differences occur in various wood characteristics, including specific gravity, mechanical properties (Bendtsen and Senft, 1984), cell length (Shiokura, 1982, Yang et al., 1986), and pulp yields (Zobel and Sprague, 1998).

The usual pattern for conifers is that wood density and MOE increase and MFA decreases as trees become older (Dungey et al. 2006). As such, juvenile wood is mostly undesirable due to its low density, low strength, high content of compression wood, high cellulose microfibril angle (MFA, the angle between the prevailing cellulose orientation and the long cell axis), low crystallinity and general high variability, as compared to mature wood (Barnett and Jeronimidis 2009; Mellerowicz et all., 2001; Zobel and van Buijtenen 1989).

MFA, is one of the key determinants of solid-wood quality due to its strong influence on the stiffness, strength, shrinkage properties and dimensional stability of structural lumber (Be0e towards the bark (Zobel and Jett 1995). Furthermore, it has been shown that in conifers, the average MFA of the S2 layer lies between 5°- 20° in mature wood (BOWYER and SHMULSKY 2007; Donaldson 2008).

The radial pith-to-bark variation of MFA is also altered by other environmental factors, whereby MFA values increase in compression wood, decrease in tension wood, and mostly increase following fertilization and thinning (Donaldson 2008).

In recent decades, most forest industries moved their attention towards usage of fast-growing tree plantations (Gräns et al. 2009), which implies that trees are harvested at younger ages than before and subsequently have greater proportion of juvenile wood (Larson 2001; Zobel and Van Buijtenen 1989). However, the negative impact of using fast-growing plantations can be reduced by changing the proportion of juvenile wood through breeding (Abdel-Gadir and Krahmer 1993; Gapare et al. 2006; Zobel and Jett 1995). In addition to selecting for trees with improved juvenile wood, it is also possible to select for an earlier transition from juvenile to mature wood in breeding programs (Gapare et al. 2006).

Transition from juvenile to mature wood usually occurs over two to five growth rings, depending on the wood property (Alteyrac et al. 2006; Mutz et al. 2004). The point at which this transition occurs is of great importance for forest managers and tree breeders as it determines the quality and value of end-use products. However, it is difficult to estimate this boundary with adequate reliability as there is usually no clear demarcation line between juvenile wood and mature wood in a tree stem (Mutz et al. 2004; Zobel and Sprague 1998). The distinction between juvenile wood and mature wood has mostly been determined by analyzing trends of radial variation (from pith to bark) for different wood properties such as density (Alteyrac et al. 2006; Mansfield et al. 2007), MOE (Wang and Stewart 2013), fiber length, and MFA (Bhat et al. 2001; Mansfield et al. 2009; Wang and Stewart 2012). This method is so-called threshold or graphic method whereby plots of each wood property are visually evaluated to locate a ring number or cambial age when the property reaches the threshold value for mature wood (Clark et al. 2007). An alternative approach is to use mathematical methods such as segmented regression (Abdel-Gadir and Krahmer 1993; Gapare et al. 2006; Szymanski and Tauer 1991) and segmented non-linear models (Alteyrac et al. 2006; Koubaa et al. 2007; Mutz et al. 2004).

It is well recognized that the proportion of juvenile wood and timing of transition in a tree is influenced by both genetic and environmental factors (Abdel-Gadir and Krahmer 1993; Mansfield et al. 2007). For instance, high heritabilities have been reported for age of transition and juvenile-wood proportion in loblolly pine *(Pinus taeda L*.) specific gravity and tracheid length (Loo et al. 1985; Stonecypher and Zobel 1966). Similarly, genetic control of time of transition has been reported to be high in slash pine (*Pinus elliottii* Engelm.) (Hodge and Purnell 1993).

The genetic control of MFA as a function of cambial age in two progeny trials of Norway spruce (*Picea abies* L.) (Chen et al. 2014) and two progeny trials of lodgepole pine (*Pinus contorta* Dougl.) (Hayatgheibi et al. 2017) have been recently investigated in Sweden. Estimated heritabilities fluctuated near the pith and then stabilized after the cambial age of 10 years in both species (Chen et al. 2014; Hayatgheibi et al. 2017). However, genetic control of MFA transition from juvenile to mature was not considered in those studies. The main objective of this study was therefore to quantify genetic variation of transition between juvenile and mature MFA based on application of different regression functions in these species, using similar progeny trials to those used by Chen et al. (2014) and Hayatgheibi et al. (2017).

## Material and Methods

### Study materials and trial design

This study utilized data from two large open-pollinated progeny trials of Norway spruce, located in southern Sweden, and two genetically unrelated open-pollinated progeny trials of lodgepole pine, located in northern Sweden (Table 1). Trials of Norway spruce, S21F9021146 aka F1146 (trial 1) and S21F9021147 aka F1147 (trial 2), comprised of 1,373 and 1,375 half-sib families, respectively, were established in 1990. Both trials were planted as randomized incomplete blocks with single-tree plots at spacing of 1.4 × 1.4 m. A set of 524 families within 112 sampling provenances were selected for this study. More detailed information about trial characteristics can be found in (Chen et al. 2014).

**Table 1:**
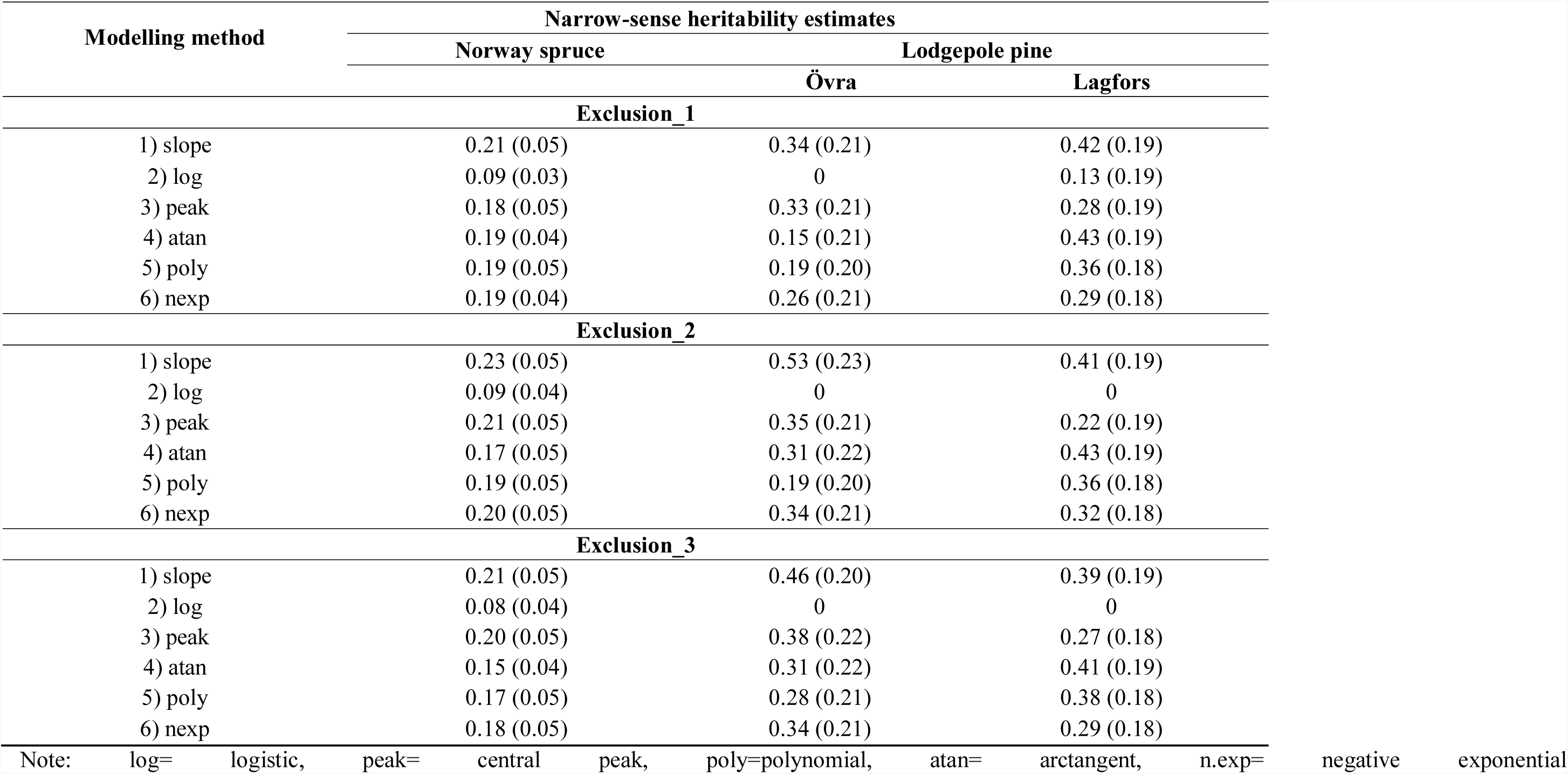
Narrow-sense heritability estimates of Norway spruce (combined-trial) and lodgepole pine (Övra and Lagfors) MFA transition from juvenile to mature wood obtained from six different regression functions applying three different exclusion methods. Standard errors of the heritabilities are in parenthesis.

| Modelling method | Narrow-sense heritability estimates |  |  |
| --- | --- | --- | --- |
|  | Norway spruce | Lodgepole pine |  |
|  |  | Övra | Lagfors |
| **Exclusion_1** |  |  |  |
| 1) slope | 0.21 (0.05) | 0.34 (0.21) | 0.42 (0.19) |
| 2) log | 0.09 (0.03) | 0 | 0.13 (0.19) |
| 3) peak | 0.18 (0.05) | 0.33 (0.21) | 0.28 (0.19) |
| 4) atan | 0.19 (0.04) | 0.15 (0.21) | 0.43 (0.19) |
| 5) poly | 0.19 (0.05) | 0.19 (0.20) | 0.36 (0.18) |
| 6) nexp | 0.19 (0.04) | 0.26 (0.21) | 0.29 (0.18) |
| **Exclusion_2** |  |  |  |
| 1) slope | 0.23 (0.05) | 0.53 (0.23) | 0.41 (0.19) |
| 2) log | 0.09 (0.04) | 0 | 0 |
| 3) peak | 0.21 (0.05) | 0.35 (0.21) | 0.22 (0.19) |
| 4) atan | 0.17 (0.05) | 0.31 (0.22) | 0.43 (0.19) |
| 5) poly | 0.19 (0.05) | 0.19 (0.20) | 0.36 (0.18) |
| 6) nexp | 0.20 (0.05) | 0.34 (0.21) | 0.32 (0.18) |
| **Exclusion_3** |  |  |  |
| 1) slope | 0.21 (0.05) | 0.46 (0.20) | 0.39 (0.19) |
| 2) log | 0.08 (0.04) | 0 | 0 |
| 3) peak | 0.20 (0.05) | 0.38 (0.22) | 0.27 (0.18) |
| 4) atan | 0.15 (0.04) | 0.31 (0.22) | 0.41 (0.19) |
| 5) poly | 0.17 (0.05) | 0.28 (0.21) | 0.38 (0.18) |
| 6) nexp | 0.18 (0.05) | 0.34 (0.21) | 0.29 (0.18) |

Note: log= logistic, peak= central peak, poly=polynomial, atan= arctangent, n.exp= negative exponential

Furthermore, two trials of lodgepole pine, Övra (Skogforsk S23F8060373) with 178 families, and Lagfors with 214 families (Skogforsk S23F7960), were established in 1980 and 1979, respectively. These families at both trials were planted in a randomized complete block (RCB) design. Each family was represented by 10 trees planted in a row with five replicates (blocks), resulting in 50 planted trees per family. Tree spacing was 2 m between rows and 1.5 m within rows. More details about trials characteristics have been further described in (Hayatgheibi et al. 2017).

### Sampling and SilviScan measurement

Increment cores (12-mm in diameter) were collected at breast height (1.3 m) from a total of 5618 Norway spruce, aged 21 years, and 823 lodgepole pine trees, aged 34-35 years, and assessed by a SilviScan instrument (Innventia, now part of RISE, Stockholm, Sweden). Before the SilviScan measurement, each increment core was sawn into a 7 mm high × 2 mm thick radial strip from the pith to the bark. The SilviScan system combines image analysis with X-ray absorption and X-ray diffraction to determine high-resolution pith-to-bark radial variations for several important wood properties, including wood density, MFA and MOE (Evans 2006; Evans and Ilic 2001). The variations in MFA from pith to bark of each core was measured as averages across consecutive 2 mm wide intervals. The annual rings were identified from the corresponding variations in wood density, and the average MFA for all rings were calculated. The number of annual rings ranged from 6 to 19 for Norway spruce and from 20 to 32 for lodgepole pine.

## Model fitting and determination of MFA transition

### Data exclusion

The radial patterns for MFA of all 5618 Norway spruce and 823 lodgepole pine individual trees were plotted against the cambial age. However, there were some individuals for which the general MFA decreasing trend from pith to the bark had been changed, due to some disturbances such as compression wood. Therefore, such outliers were identified and excluded prior to data analysis. Following removal of such outliers, the remaining Norway spruce and lodgepole pine individuals having fewer than 12 and 20 annual rings, respectively, were also excluded from data analysis. The three applied exclusion methods are as follows:

1. Exclusion_1 or basic method: exclusion of those individuals for which MFA values increased with cambial age (abnormal curves). This was the baseline of data treatment and thus the first step of exclusion methods 2 and 3. This first exclusion step resulted in exclusion of about 2.8 % and 5.1 % of Norway spruce (Höreda and Erikstorp, respectively) and 0.2 % and 1.7 % of lodgepole pine individuals (Lagfors and Övra, respectively)
2. Exclusion_2 or shape-based method: following the baseline method, annual rings of the individuals for which average MFA values were greater than their three previous rings average MFA, were removed. Based on this method, about 480 and 959 annual rings in Norway spruce (Höreda and Erikstorp, respectively) and about 613 and 398 annual rings in lodgepole pine (Lagfors and Övra, respectively) were removed. This resulted in removal of about 4.0 % and 7.5 % of Norway spruce (Höreda and Erikstorp, respectively) and 0.9 % and 7.3 % of lodgepole pine individuals (Lagfors and Övra, respectively).
3. Exclusion_3 or family-based method: the data after exclusion with the baseline method were exposed to another method of data exclusion. Annual rings, which had average MFA values deviating from their corresponding family-mean MFA values by more than 1.96 × SD, were excluded from data analysis. Based on this method, about 838 and 665 annual rings in Norway spruce (Höreda and Erikstorp, respectively) and about 177 and 159 annual rings in lodgepole pine (Lagfors and Övra, respectively) were removed. This method resulted in the removal of about 3.4 % and 5.2 % of Norway spruce (Höreda and Erikstorp, respectively) and 0.2 % and 1.5 % of lodgepole pine individuals (Lagfors and Övra, respectively).

### Regression functions and MFA transition

After removal of outliers, six different regression functions were fitted to the pith-to-bark MFA profiles of the individual trees. A threshold value of 20° was considered for MFA and when the parameter of the fitted functions fell below the threshold, the estimated parameter was defined as MFA transition.

All data analysis was carried out using the R statistical programming environment (R Development Core).

The fitted models were as follow:

1. *Slope function*

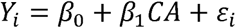

where *β*_0_ is the intercept and *β*_1_ is the slope. In all equations, *Y*_i_ is the MFA value of the tree *i*, is the cambial age, and *ε*_i_ is the random error.
2. *Logistic function*

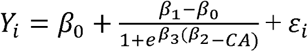

where *β*_0_ is the upper asymptote MFA, *β*_1_ is the lower asymptote MFA, *β*_2_ is the inflection point, and *β*_3_ is the sharpness.
3. *Central peak*

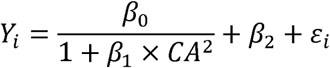

where *β*_0_ + *β*_2_ is the upper asymptote MFA, 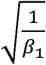 is the inflection point, and *β*_2_ is the lower asymptote MFA.
4. *Polynomial* The radial profile of MFA from each tree was plotted with respect to cambial age and fitted with a third-order polynomial regression as below:

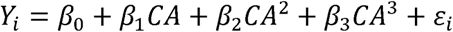
5. *Arctangent*

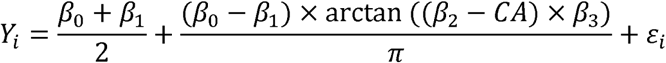

where *β*_0_ is the upper asymptote MFA, *β*_1_ is the lower asymptote MFA, *β*_2_ is the inflection point, and *β*_3_ is the slope.
6. *Negative exponential*

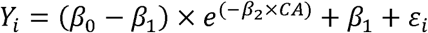

where *β*_0_ is the MFA at cambial age zero, *β*_1_ is the lower asymptote MFA, and *β*_2_ is the slope at cambial age zero.

### Genetic analysis

Variance components for genetic analysis were estimated using ASReml 3.0 (Gilmour et al. 2009) based on linear mixed-effects model using joint-site analysis for Norway spruce and single-site analysis for lodgepole pine as follows:

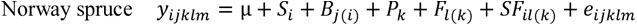

where *y*_*ijklm*_ is the observation on the *m*th tree from the *l*th family within the *k*th provenance in the *l*th block within the *i*th site. µ is the overall mean; *S*_*i*_ is the fixed effect of the *i*th site; *B*_*j(i)*_ is the fixed effect of the *j*th block within the *i*th site; *P*_*k*_ is the fixed effect of the *k*th provenance; *F*_*l(k)*_ is the random effect of the *l*th family within the *K*th provenance; *SF*_*il(k)*_ is the random interaction effect of the *i*th site and the *l*th family within the *k*th provenance; and *e*_*ijklm*_ is the random residual effect.

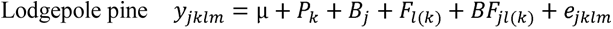

where *y*_*jklm*_ is the vector of observation on the *m*th tree from the *l*th family within the *k*th provenance in the *j*th block. µ is the overall mean; *P*_*k*_ and *S*_*j*_ are the fixed effects of the *k*th provenance and the *j*th block, respectively; *F*_*l(k)*_ is the random effect of the *l*th family within the *k*th provenance; *BF*_*jl(k)*_is the random interactive effect of the *j*th block and the *l*th family within the *k*th provenance and *e*_*jklm*_ is the random residual effect.

Estimates of heritability were obtained using variance components from the univariate join-site (for Norway spruce) and single-site (for lodgepole pine) analysis. Approximate standard errors were estimated using the Taylor series expansion method (Gilmour et al. 2009). Individual-tree narrow-sense heritability 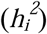 for MFA of each mathematical model was calculated using the following equations assuming these open-pollinated family are half-sib families (Falconer and Mackay 1996):

Norway spruce

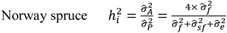

Lodgepole pine

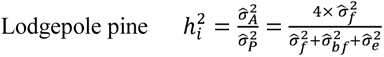

where 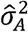 is the additive genetic variance; 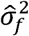 is among family variance; 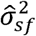 is the site by family variance; 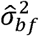 is the family by block variance; and 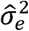 is the residual variance.

Pooled-site analysis genetic gain (*∆G*_*t*_), expressed as percentage in direct selection of MFA transition-age was estimated as:

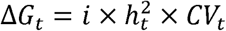

where *i* is the selection intensity of 1% (*i*=2.67), 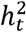 is the narrow-sense heritability of MFA transition and *CV* is the coefficient of variation of the phenotypic effect (calculated as the phenotypic standard deviation divided by the mean of the trait).

## Results

### MFA transition and model fitting in Norway spruce and lodgepole pine MFA radial variation

For both species, the MFA average profiles decreased from high values close to the pith towards stable levels close to the bark (Fig. 1). In Norway spruce, mean MFA profile decreased from about 30° at the pith and then stabilized after reaching a cambial age of 10 years at about 10° in Höreda and about 12° in Erikstorp. The mean MFA profile for lodgepole pine started at 40° followed by a rapid decrease to about 30° after 3 years, after which the shape of the development was similar to that for Norway spruce but slower, stabilizing after cambial age of 15 years at about 12° in Övra and about 10° in Lagfors.

**Fig. 1.**
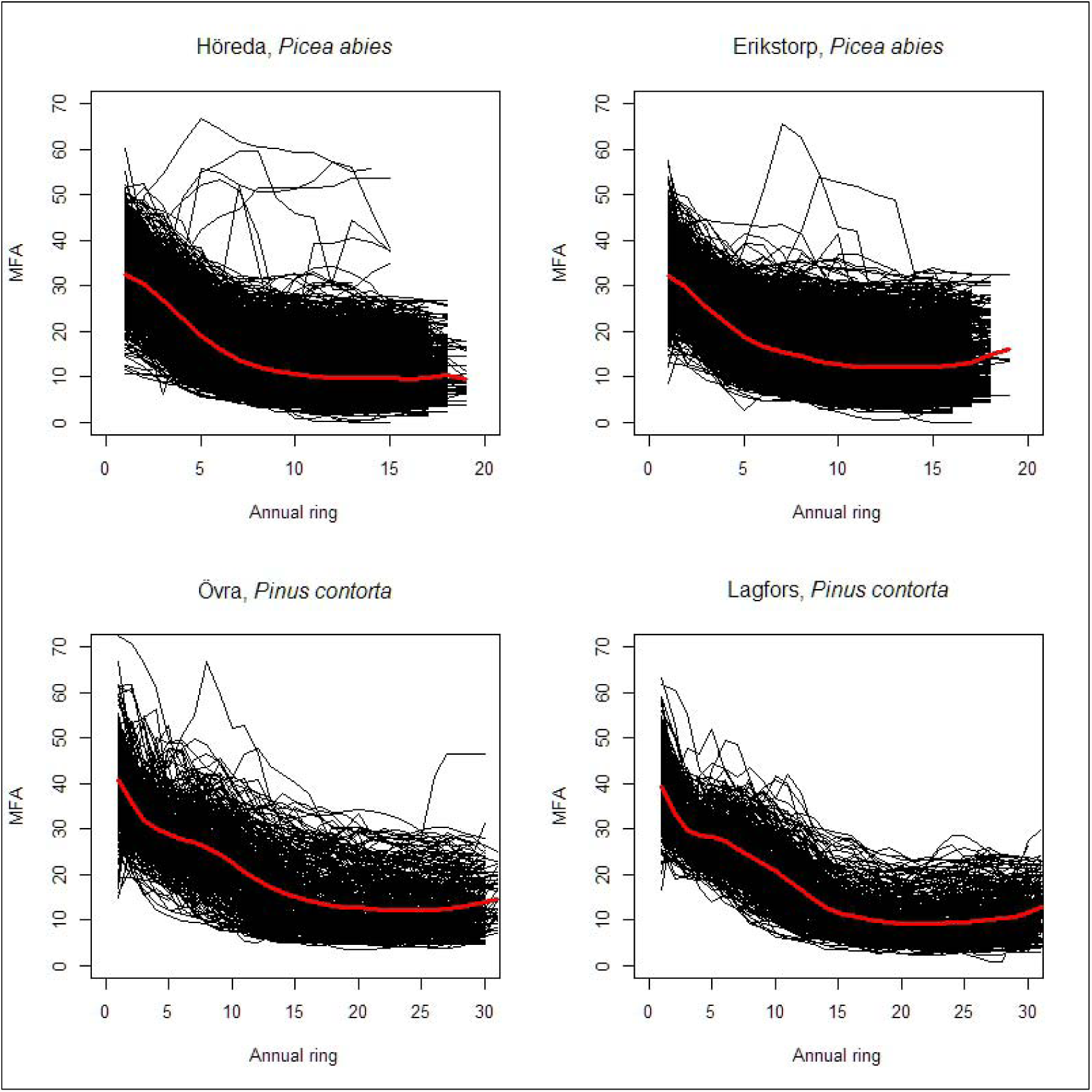
Radial trends for MFA of *Picea abies* at two trials (Höreda and Erikstorp) from cambial age 1 to 20 and for *Pinus contorta* at two trials (Övra and Lagfors) from cambial age 1 to 30. The black lines represent the actual observations from all individual trees and the red line is the mean radial variation of MFA against the cambial age.

### Model fitting and MFA transition

For each fitted regression function, different threshold values were tested to define the best model fit for individual trees transition from juvenile to mature MFA. Although the threshold value of 20° gave the best model fit, there were still some individuals for which the MFA profile never passed the threshold, thus, they remained constantly above or below the threshold. Figures 2 to 5 show the proportion of Norway spruce and lodgepole pine individuals for which the MFA transition was estimated based on the applied six different regression functions, under the threshold of 20°.

**Fig. 2.**
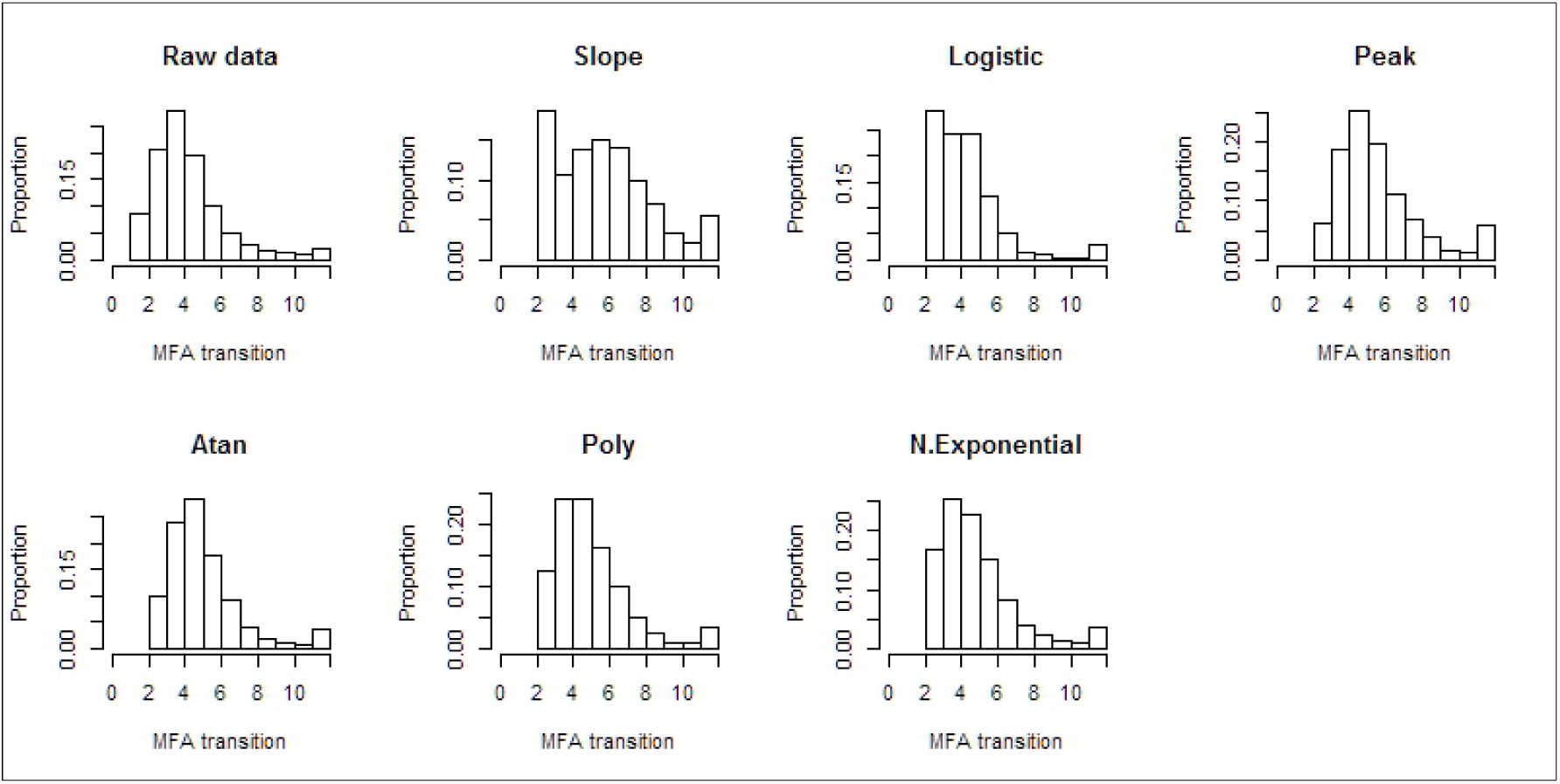
Proportion of Norway spruce (Höreda) transition age under the mean (raw data) and six different regression functions using the threshold level of 20 degrees for MFA.

**Fig. 3.**
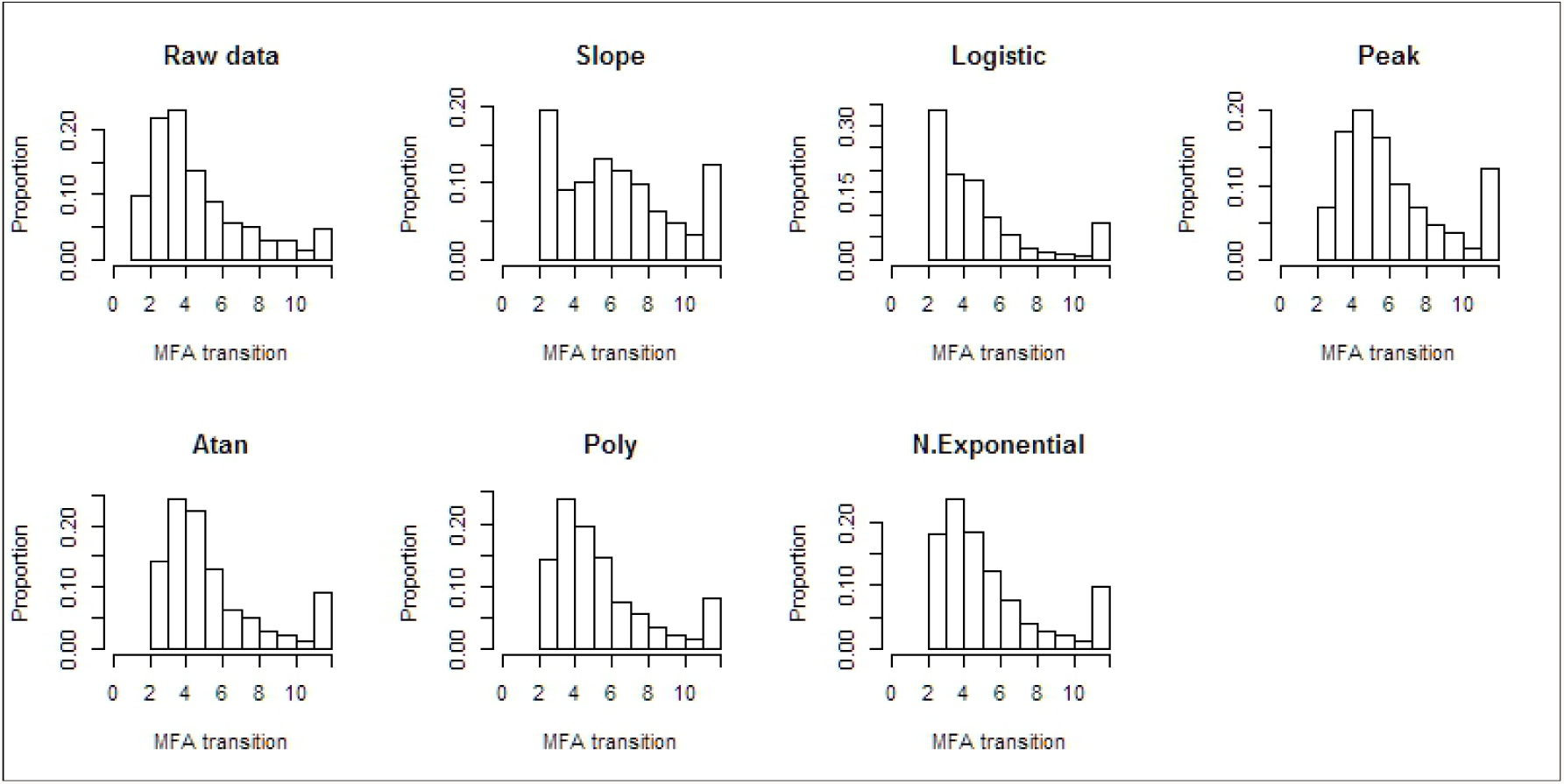
Proportion of Norway spruce (Erikstorp) transition age under the mean (raw data) and six different regression functions using the threshold level of 20 degrees for MFA

### Heritability estimates

Narrow-sense heritability estimates were obtained for combined-site Norway spruce and single-site lodgepole pine MFA transition.

**Fig. 4.**
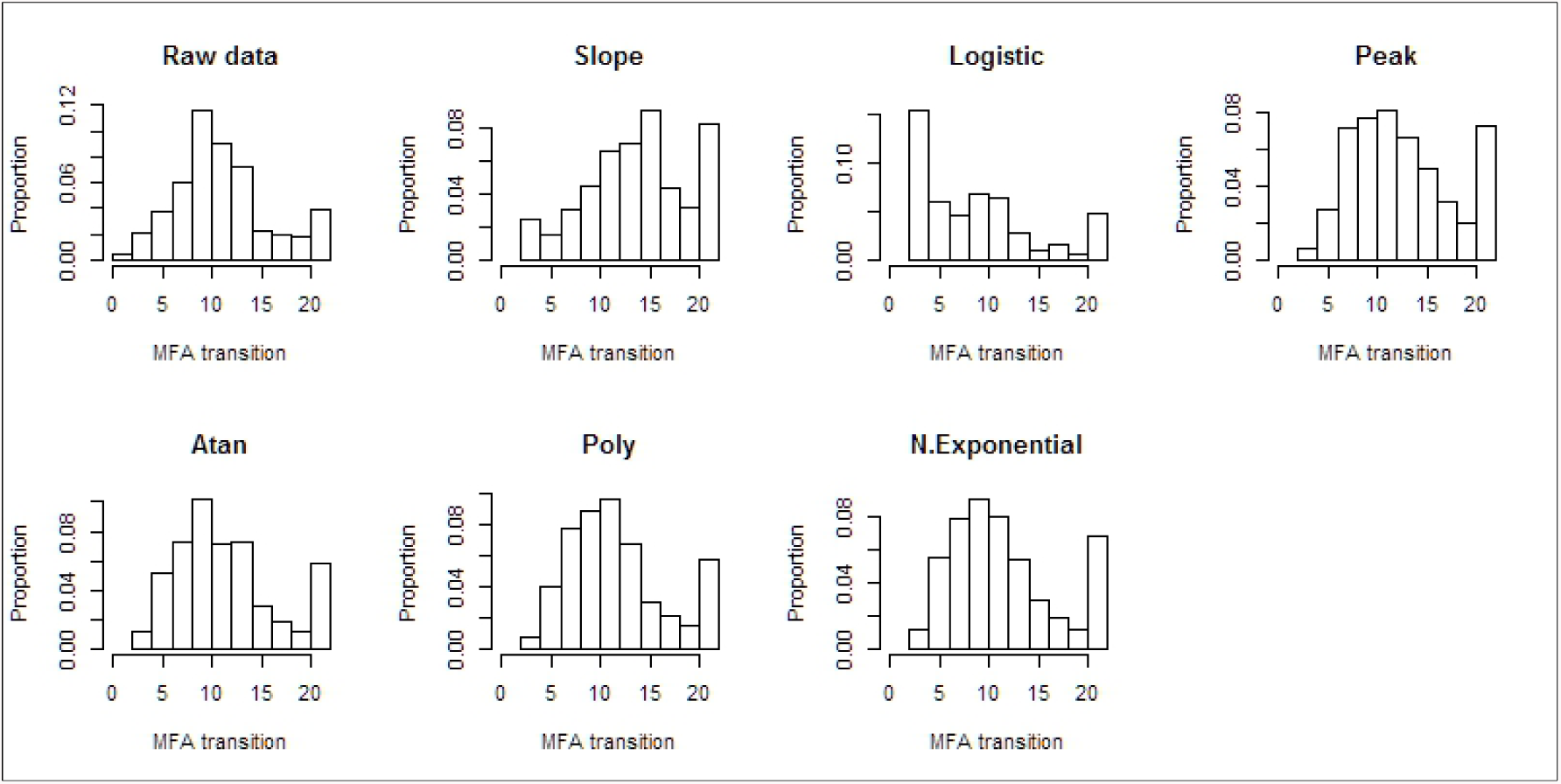
Proportion of lodgepole pine (Övra) transition age under the mean (raw data) and six different regression functions using the threshold level of 20 degrees for MFA

### Norway spruce

Estimated heritabilities were mostly stable following data exclusions. However, heritabilities were slightly higher after shape-based exclusion (exclusion_2), while they were slightly lower after the family-based exclusion (exclusion_3), particularly for arctangent, polynomial and negative-exponential functions (Table 1). In general, *h*^2^ ranged from 0.08 to 0.23 with the highest estimate obtained based on the slope function and the lowest estimate obtained based on the logistic function, under all exclusion methods.

**Fig. 5.**
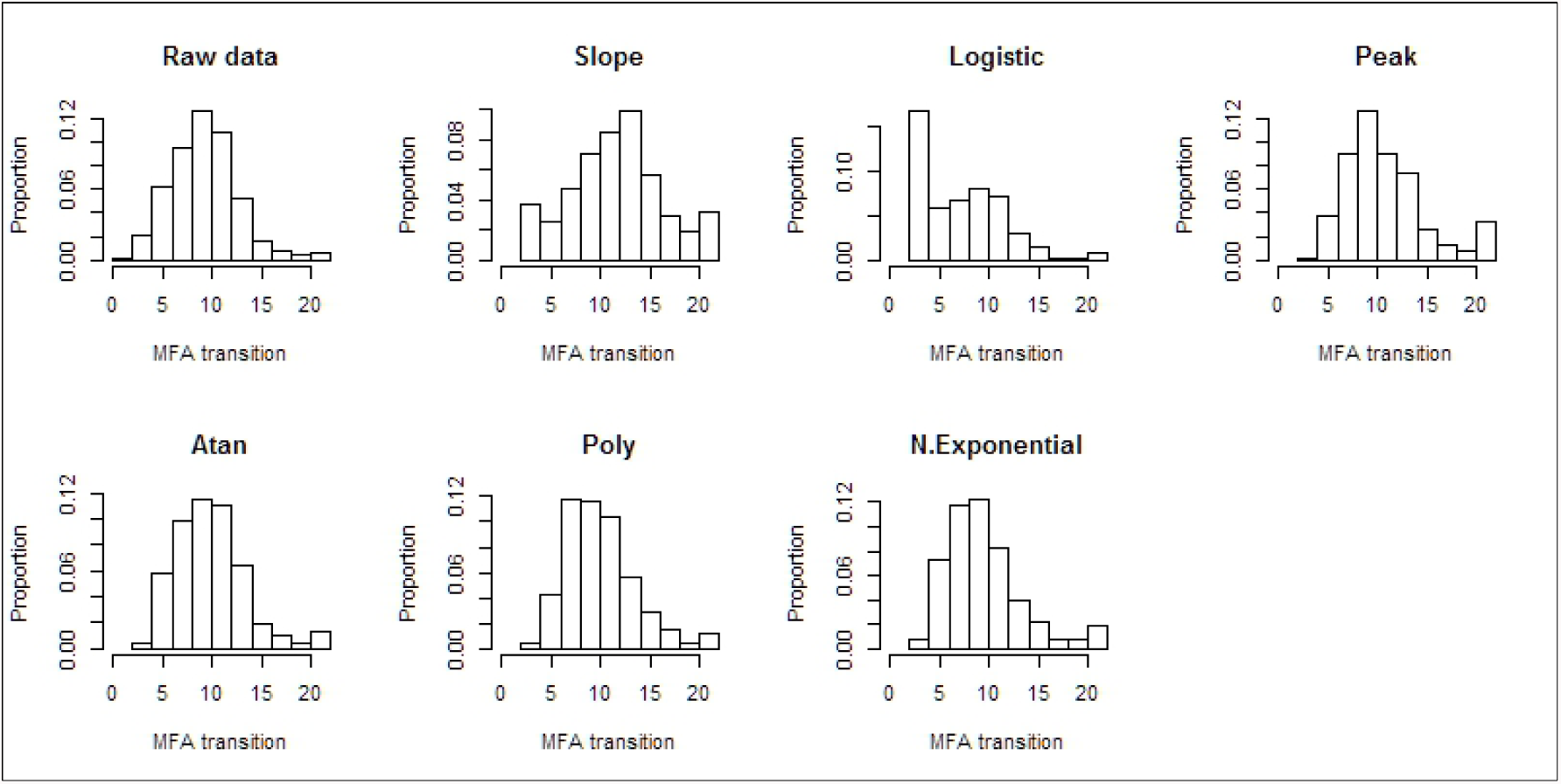
Proportion of lodgepole pine (Lagfors) transition age under the mean (raw data) and six different regression functions using the threshold level of 20 degrees for MFA

### Lodgepole pine

In general, *h*^2^ estimates observed for lodgepole pine were higher than those for Norway spruce (Table 1). Heritabilities in Övra increased significantly after exclusion_2 and exclusion_3, particularly after exclusion_2, while they slightly decreased after exclusion_3 in Lagfors. As similarly observed for Norway spruce, heritability estimates were lowest based on the logistic function, while they were highest based on the slope function. In general, heritabilities ranged from 0.15 to 0.53 in Övra and from 0.22 to 0.43 in Lagfors (logistic function was excluded). In addition to the slope function, high heritabilities obtained based on the central peak (ranging from 0.33 to 0.38) and negative exponential functions (ranging from 0.26 to 0.34) in Övra, and based on the arctangent (ranging from 0.41 to 0.43) and polynomial (ranging from 0.36 to 0.38) in Lagfors.

### Genetic gain

As was observed for heritability estimates, genetic gains obtained based on the slope function were highest, while those obtained based on the logistic function were lowest, regardless of which exclusion method was applied (Table 2). High genetic gains were also obtained based on the negative-exponential function for both species when using exclusion-1. Similarly, application of exclusion-2 and exclusion-3 led to high genetic gains when MFA was modelled by negative-exponential and central peak functions in Norway spruce and by arctangent and negative-exponential functions in lodgepole pine (Table 2).

**Table 2.** Genetic gains (∆*G*_*t*_ %) obtained for transition age in combined-site Norway spruce and combined-site lodgepole pine using the selection intensity of 1%

| Norway spruce |  |  |  |
| --- | --- | --- | --- |
| Model | $\Delta G_t$ (%) | | |
|  | exclusion-1 | exclusion-2 | exclusion-3 |
| Slope | 27.4 | 29.4 | 27.1 |
| Log | 13.3 | 13.0 | 11.1 |
| Peak | 20.9 | 24.1 | 23.2 |
| Atan | 23.2 | 20.2 | 17.1 |
| Poly | 23.3 | 23.0 | 21.6 |
| N. ex | 25.0 | 25.7 | 20.1 |

| lodgepole pine |  |  |  |
| --- | --- | --- | --- |
| Model | $\Delta G_t$ (%) | | |
|  | exclusion-1 | exclusion-2 | exclusion-3 |
| Slope | 40.2 | 48.4 | 45 |
| Log | 0 | 3.7 | 11 |
| Peak | 32.8 | 30.2 | 35.7 |
| Atan | 29.4 | 39.5 | 39.9 |
| Poly | 29.6 | 29.7 | 35.6 |
| N. ex | 33.4 | 38.2 | 38.9 |

Note: Log= logistic, Peak= Central peak, Poly=polynomial, Atan= arctangent, N.exp= negative exponential

## Discussion and conclusion

Several investigations in various conifers have determined the age for MFA transition from juvenile to mature, using different linear and non-linear models (Clark et al. 2007; Mansfield et al. 2009; Wang and Stewart 2012). However, no study to date has examined genetic variation of MFA transition and its possible genetic gain based on employing different regression functions. Few studies have investigated the genetic control of transition age in fast growing pines (Gapare et al. 2006). As such, a family heritability estimate of 0.36 and moderate genetic gains in selection of early age of transition for specific gravity in loblolly pine has been reported (Loo et al. 1985). Similarly, moderate genetic gains were reported for latewood density transition age in radiata pine (*Pinus radiata* D. Don) (Gapare et al. 2006).

One of the central goals in forest tree breeding is to reduce costs and time of breeding cycles by making genetic gains as early as possible. Additionally, due to the increased demand in fast-grown plantations, which contain more juvenile wood, it is of vital importance to breed for more uniform wood, or a wood with lower proportion of juvenile wood.

Therefore, the main focus of this study was to evaluate the degree of genetic control in MFA transition from juvenile to mature in Norway spruce and lodgepole pine to ensure that future end-use products obtained from these species have desirable wood characteristics.

Although the MFA average profiles started to stabilize at about 15° in both species (Fig. 1), the chosen threshold value of 20° resulted in the best model fit for estimation of individual trees MFA transition from juvenile to mature. Furthermore, as the wood of conifers having MFA of 20° is considered as mature (Bowyer and Shmulsky 2007; Donaldson 2008), such a high transition degree for MFA will enables incorporation of wood quality traits into selective breeding programs of Norway spruce and lodgepole pine as early as possible.

Most of the narrow-sense heritability estimates obtained in this study were statistically significant, except for those obtained based on the logistic function under all exclusion models. This might be driven by the inability of this function to estimate MFA transition under the selected threshold, particularly in lodgepole pine.

Overall, heritability estimates obtained for lodgepole pine were generally greater than those obtained for Norway spruce. This is in line with results of the studies by Chen et al. (2014) and Hayatgheibi et al. (2017), as *h*^2^ obtained for area-weighted MFA in two progeny trials of lodgepole pine were greater than that obtained for Norway spruce. Similarly, heritability estimates obtained based on the slope function were highest in both species across all exclusion methods. However, *h*^2^ obtained for Norway spruce based on negative exponential and central peak, were also high. Similarly, high *h*^2^ were observed based on the negative exponential and central peak functions in Övra, and based on the arctangent and polynomial functions in Lagfors.

Results of this study, which made use of large genetic data sets of two important coniferous species, revealed that there is a possibility to breed for earlier MFA transition from juvenile to mature as genetic gains achieved for this trait were very high.

In general, genetic gains obtained for MFA transition, except for those obtained based on the logistic function, were higher in lodgepole pine due to the higher narrow-sense heritability estimates obtained for this trait in lodgepole pine. Further, in addition to the slope function, highest genetic gains were achieved based on the negative exponential and central peak functions in Norway spruce when data trimming by exclusion-2 was applied, and based on the negative exponential and arctangent functions in lodgepole pine after applying exclusion-3. Findings of this study indicate that there is a possibility to select for a reduction in MFA transition age, and therefore, a decrease in proportion of the log containing juvenile wood in Norway spruce and lodgepole pine selective breeding programs.

## Acknowledgements

The authors gratefully acknowledge financial support from Föreningen Skogsträdsförädling, Bo Rydins, Kempe foundations, and Swedish University of Agricultural Sciences (SLU). The samples were collected in Skogforsk’s operative breeding experiments. We also acknowledge Anders Fries who collected lodgepole pine increment cores, Åke Hansson, Thomas Trost, Lars Olsson and Thomas Grahn for preparation of the cores, analyses with SilviScan and preparation of the trait data, as well as Johan Kroon who prepared detailed lodgepole pine genetic data. We also want to acknowledge Bio4Energy for their contribution to the SilviScan analysis of Norway spruce data to the Knut and Alice Wallenberg’s foundation that provided support for Norway spruce samples and data collection.

